# Relapse to cotinine seeking in rats: Differential effects of sex

**DOI:** 10.1101/2021.10.14.464393

**Authors:** Xiaoying Tan, Elizabeth Neslund, Zheng-Ming Ding

## Abstract

Relapse is a defining feature of smoking and a significant challenge in cessation management. Elucidation of novel mechanisms underlying relapse may inform future treatments. Cotinine, the major metabolite of nicotine, has been shown to support intravenous self-administration in rats, suggesting it as one potential mechanism contributing to nicotine reinforcement. However, it remains unknown whether cotinine would induce relapse-like behaviors. The current study investigated relapse to cotinine seeking in two relapse models, the reinstatement of drug seeking and incubation of drug craving models. In the reinstatement model, rats were trained to self-administer cotinine, extinguished cotinine-associated responses, and underwent cue-, drug-, or stress-induced reinstatement. Conditioned cues associated with cotinine self-administration, cotinine (1-2 mg/kg), or the pharmacological stressor yohimbine (1.25-2.5 mg/kg) reinstated cotinine seeking. Female rats displayed more pronounced cue-induced, but not drug- or stress-induced reinstatement than male rats. In addition, an overall analysis revealed that female rats exhibited greater cotinine self-administration, but less extinction than male rats. In the incubation model, rats were trained to self-administer cotinine, and underwent forced withdrawal in home cages. Rats were tested for cue-induced cotinine seeking on both withdrawal day 1 and withdrawal day 18. Rats exhibited greater cotinine-seeking on withdrawal day 18 compared to withdrawal day 1, with no difference between male and female rats. These findings indicate that cotinine induces sex-dependent relapse to cotinine seeking in rats, suggesting that cotinine may be a novel mechanism contributing to relapse. These rat models are valuable preclinical tools for interrogation of neurobiological underpinnings of relapse to cotinine seeking.

**Significance Statement:** Relapse is a defining feature of smoking and a significant challenge in cessation management. Elucidation of novel mechanisms underlying relapse may inform future treatment. Cotinine, the major metabolite of nicotine, has previously been shown as a potential mechanism underlying the development of nicotine reinforcement because it supports intravenous self-administration in rats. The current study found that conditioned cues, priming injection of cotinine, and acute exposure to the pharmacological stressor yohimbine induced robust cotinine-seeking behavior in rats with a history of cotinine self-administration. In addition, significant sex effects were revealed in that female rats exhibited greater cotinine self-administration, less extinction, and more robust cue-induced reinstatement of cotinine seeking than male rats. These findings suggest that cotinine may be a novel mechanism contributing to relapse to nicotine seeking. In addition, these rat models are valuable preclinical tools for interrogation of behavioral and neurobiological underpinnings to relapse to cotinine seeking, thus forming basis for developing effective therapeutic strategy to aid in smoking cessation.

## Introduction

Cigarette smoking continues to be a leading public health concern and cause of preventable death in the United States (U.S. Department of Health and Human Services (HHS) 2014). Smoking rate remained at approximately 17% in Americans age 12 and above in 2018 (Substance Abuse and Mental Health Services Administration (SAMHSA) 2019). Recent surge of electronic cigarette use has resulted in estimated 3.6 million users in middle and high school students in the US in 2020, posing additional risk to youth (Wang et al 2020a). Nicotine is the major addictive component in cigarettes, and it activates brain nicotinic acetylcholine receptors (nAChRs) to produce its effects (Benowitz 2010). Several FDA-approved medications (e.g., nicotine replacement therapy and varenicline) mainly target nicotine’s effects to aid in smoking cessation in clinics. However, these medications only provide modest clinical benefits (Aubin et al 2014, Prochaska & Benowitz 2016).

Relapse to smoking after a period of abstinence is common in smokers and a major obstacle in smoking cessation management. Most smokers attempt to quit smoking, but encounter high rates of failure. For example, in 2018, approximately 55% of adult smokers made quit attempts, but only 7.5% succeeded (Creamer et al 2019). Most quit attempts are unassisted with medications or behavioral interventions, and most relapse occurs within the first week after a given quit attempt with only 2-5% achieving long-term abstinence for 6-12 months (Hughes et al 2004, Prochaska & Benowitz 2016). Even with medication-assisted smoking cessation, the abstinence rates are modest. A meta-analysis of randomized controlled trials of three FDA-approved medications revealed that fewer than 40% of smokers achieved abstinence at 3 months following treatments, and benefits quickly diminished with quit rates dropping to below 20% at 12 months (Rosen et al 2018). Therefore, there is critical need for further understanding of relapse mechanisms to inform future treatments.

In humans with substance use disorders, relapse is typically precipitated by risk factors such as stress, drug-related cues, and the drug itself (Doherty et al 1995). Decades of preclinical research has developed and validated animal models that capture these characteristics of relapse (Venniro et al 2016). The reinstatement of drug seeking model is one of the most studied models. This is an extinction-based model in which animals are first trained to self-administer a drug, then undergo extinction training to extinguish drug-maintained behavior followed by reinstatement of previously drug-reinforced behavior upon acute exposure to the drug, drug-associated cues, or a stressor (Shaham et al 2003). Animals trained to self-administer nicotine demonstrated robust reinstatement of nicotine seeking following non-contingent administration of small doses of nicotine (Chiamulera et al 1996), contingent presentation of environment cues previously associated with nicotine self-administration (Forget et al 2009), and acute exposure to either pharmacological or physical stressors, e.g., the α2 adrenergic receptor antagonist yohimbine (Grella et al 2014) or electrical foot shock (Buczek et al 1999). Another influential model is the incubation of drug craving model that is an abstinence-based relapse model. In this model, animals are first trained to self-administer a drug, undergo a period of abstinence or withdrawal period in the home cage with no drug exposure, and then are tested, at various time points, in the drug self-administration environment with the presentation of only drug-associated cues. Time-dependent increase in cue-induced drug seeking indicates the incubation of craving (Pickens et al 2011). Animals self-administering nicotine showed increased nicotine seeking in this model (Funk et al 2016, Markou et al 2016). All these models have been successful used to reveal important mechanisms underlying relapse to nicotine seeking in animals (Stoker & Markou 2015).

Cotinine is a minor tobacco alkaloid and the major metabolite of nicotine. Approximately 70- 80% of absorbed nicotine is converted to cotinine through a hepatic enzymatic process (Benowitz & Jacob 1994). It is commonly used as a biomarker for nicotine exposure due mainly to its long plasma half-life (Hukkanen et al 2005). Cotinine penetrates the blood brain barrier and produces various neurobehavioral effects (Tan et al 2021). There is evidence suggesting that cotinine can act as a weak agonist of nAChRs with its potency being several orders of magnitude less than nicotine (Abood et al 1983, Anderson & Arneric 1994). Cotinine is a neuroactive metabolite of nicotine and produces its own effects. In animals, cotinine altered food-reinforced behaviors (Goldberg et al 1989, Risner et al 1985), changed neurotransmission in the brain and periphery (Dwoskin et al 1999, Schroff et al 2000), and was substituted for nicotine in producing nicotine-like discriminative stimulus effects (Rosecrans & Chance 1977, Takada et al 1989). Low doses of cotinine altered locomotor activity (Wang et al 2020b, Wiley et al 2015), and modulated the effects of nicotine or alcohol on locomtion (Clemens et al 2009, Dar et al 1993). Recent studies indicate that cotinine enhanced attention, learning and memory in animal models of cognitive impairments and was beneficial in animal models of Alzheimer’s disease and schizophrenia (Boiangiu et al 2020, Echeverria et al 2011, Terry Jr et al 2005). In humans, cotinine altered subjective states related to acute smoking abstinence and withdrawal (Benowitz et al 1983, Keenan et al 1994), and interacted with nicotine patch to modulate smoking-withdrawal symptoms (Hatsukami et al 1998).

A recent study demonstrated that cotinine supported intravenous self-administration in rats. In this study, cotinine at doses producing blood cotinine concentrations consistent with levels seen in habitual smokers was self-administered in both fixed- and progressive-ratio schedules. Cotinine self-administration exhibits similarities and differences to nicotine self-administration in a dose- and schedule-dependent manner. In general, cotinine was less robust than nicotine in inducing self-administration. In addition, no difference in cotinine self-administration was observed between male and female rats (Ding et al 2021). These results suggest that cotinine may be reinforcing by itself and play a role in nicotine self-administration. However, it remained undetermined whether cotinine would induce relapse-like behavior in rats. Therefore, the current study aimed to characterize relaspe to cotinine-seeking behavior in both male and female rats using two relapse models, i.e., the reinstatement of drug seeking and incubation of craving models. Given previous findings, the hypothesis to be tested was that cotinine would induce relapse-like behavior in both male and female rats with no significant sex difference.

## Materials and Methods

### Animals

Young adult male and female (starting at ∼ 7-8 weeks old) Wistar rats (Envigo, Indianapolis, IN USA) were housed in groups (2-4/cage) upon arrival on a reversed 12-hr light-dark cycle in rooms controlled for temperature and humidity. Rats were acclimated for at least 5 days before catheterization surgery and were individually housed after surgery in cages enriched with a polycarbonate play tunnel and nestlets. Food and water were available *ad libitum* except during behavioral testing. Experiments were performed during the dark phase. Protocols used were approved by the Institutional Animal Care and Use Committee at the Pennsylvania State University College of Medicine. All experiments were performed in accordance with the principles outlined in the Guide for the Care and Use of Laboratory Animals (National Research Council 2011).

### Intravenous catheterization

Rats were implanted with a catheter into the jugular vein following a procedure detailed previously (Berg et al 2014, Ding et al 2021). Briefly, rats were anesthetized with 2-3% isoflurane inhalation and a polyurethane tubing (I.D. x O.D. = 0.63 × 1.02mm; Instech Laboratories, Inc., Plymouth Meeting, PA, USA) was inserted into the right jugular vein. The remaining portion of the catheter coursed subcutaneously over the shoulder blade to exit the back of the rat via a 22-gauge cannula (Plastics One, Roanoke, VA, USA). Bupivacaine hydrochloride (Hospira, Inc., Lake Forest, IL, USA) at 0.25% and carprofen at 5 mg/kg (Zoetis Inc., Kalamazoo, MI, USA) were applied as analgesia during surgery. Catheters were flushed daily with ∼0.5ml heparinized saline (20 IU/ml, Hospira, Inc., Lake Forest, IL, USA) containing 0.13 mg/ml gentamicin sulfate (McKesson, Livonia, MI, USA). Rats were checked once a week, typically after the Friday session, for catheter patency with intravenous administration of ∼0.1ml of 10 mg/ml methohexital sodium (Par Pharmaceutical, Chestnut Ridge, NY, USA). Rats with failed catheters were excluded from further experiment and analysis.

### Intravenous self-administration of cotinine

Self-administration was conducted in standard chambers equipped with two levers, a cue light, and a house light (Med Associates Inc., St. Albans, VT, USA), following procedures previously described (Ding et al 2021). A light food restriction procedure was implemented to maintain rats at ∼85% body weight (with 2–3 full bricks/day of standard rat chow immediately following the completion of a session). A piece of Froot Loops cereal was placed on the active lever as a bait during the first two sessions to promote exploratory behavior. A passive infusion was delivered by the experimenter at the beginning of each session. A fixed-ratio (FR) 1 schedule was used, and responses on the active lever caused an intravenous infusion of (-)-cotinine (Sigma, St. Louis, MO, USA) at 0.03 mg/kg/infusion, a dose inducing robust self-administration behavior in rats as demonstrated in a previous study (Ding et al 2021). Infusions were delivered in 55 µl over a 3s period via a syringe pump (PHM-100, Med Associates Inc., St. Albans, VT, USA). During the infusion, the house light was turned off and the cue light above the active lever was turned on. The infusion was followed by a 17s time-out period during which both the cue light and the house light were turned off. Lever presses during the infusion and time-out periods were recorded but produced no further infusions. Responses on the inactive lever were recorded, but no programmed consequences ensued. Sessions were 2 hours in duration, and were conducted daily during weekdays for 20 sessions.

### Extinction and reinstatement of cotinine seeking

Following self-administration, rats underwent daily extinction sessions for 10 sessions. During extinction, responses on neither lever led to the delivery of cotinine or light cues. Reinstatement was conducted following the conclusion of extinction. For cue-induced reinstatement, responses on the previous active lever led to contingent delivery of light cues associated with cotinine self-administration with no cotinine delivered. Responses on the previous inactive lever resulted in no programmed consequence. Drug- and stress-induced reinstatement sessions were conducted under the condition similar to extinction, i.e., no cotinine or light cues was available. For drug-induced reinstatement, rats received pretreatment of either saline or cotinine (1.0 or 2.0 mg/kg, s.c.). For stress-induced reinstatement, rats received pretreatment of saline or yohimbine (1.25, or 2.5 mg/kg, s.c.). Pretreatment was given ∼ 30 min prior to the start of a reinstatement session, and a within-subject design was employed with different doses being administered in a random order. Additional extinction sessions were included between treatment sessions to allow responses to return to baseline before the next treatment.

### Incubation of craving to cotinine-seeking behavior

This experiment followed a previously described procedure to assess cue-induced cotinine-seeking behavior following self-administration and a withdrawal period. A within-subject assessment was applied (Markou et al 2016). Briefly, rats followed the afore-mentioned procedure for self-administration of cotinine at 0.03 mg/kg/infusion on a FR1 schedule for 14 sessions. During the first day of withdrawal, rats were tested for cue-induced cotinine seeking in a session when responses on active lever resulted in the delivery of cue lights only but not cotinine. This was the ‘withdrawal day 1’ session. Rats then returned to home cages and continued forced abstinence during withdrawal days 2-17 with no access to cotinine or operant chambers. On withdrawal day 18, rats were placed back in operant chambers for another test session of cue-induced cotinine seeking with the same condition as the ‘withdrawal day 1’ session. This is the ‘withdrawal day 18’ session. The responses during the ‘withdrawal day 18’ were compared to those during the ‘withdrawal day 1’ for the determination of the incubation of cotinine craving.

### Statistical analysis

Data were expressed as mean ± SEM, and were analyzed with mixed ANOVAs with repeated measures on session followed by multiple comparisons with the Bonferroni correction. The significance level was set at *p* < 0.05.

## Results

### Cue-induced reinstatement of drug-seeking behavior

During self-administration (Fig. 1B), there was significant effect of session (F_19, 285_ = 7.8, *p* = 0.000), but not sex (F_1, 15_ = 0.8, p = 0.39) or interaction between sex and session (F_19, 285_ = 0.9, *p* = 0.53). Both male and female rats increased infusions overtime. During extinction (Fig. 1C), there were significant effects of session (F_10, 150_ = 6.2, *p* = 0.000), sex (F_1, 15_ = 13.0, *p* = 0.003), and sex x session interaction (F_10, 150_ = 4.6, *p* = 0.000). Male rats readily extinguished lever responses. For female rats, active responses increased transiently during the first three extinction sessions, and then gradually decreased during the remaining extinction sessions. This brief increase is reminiscent of an ‘extinction burst’ observed in studies with other drugs of abuse, including nicotine and cocaine (Plaza-Zabala et al 2010, Ward et al 2009). Further analysis revealed that female rats exhibited greater active responses than females during the first several extinction sessions. During reinstatement, there were significant effects of session (F_1, 15_ = 63.7, *p* = 0.000), sex (F_1, 15_ = 12.1, *p* = 0.003), and sex x session interaction (F_1, 15_ = 12.1, *p* = 0.003) on active responses (Fig. 1D, upper panel). The contingent presentation of light cues associated with cotinine self-administration increased active responses compared to extinction baseline averaged from the last three extinction session, and female rats exhibited more robust increase than male rats. On the other hand, cue presentation did not alter inactive responses (Fig. 1D, bottom panel) with no significant effect of session (F_1, 15_ = 3.1, *p* = 0.1), sex (F_1, 15_ = 1.1, *p* = 0.32), or sex x session interaction (F_1, 15_ = 1.1, *p* = 0.32).

**Figure 1.**
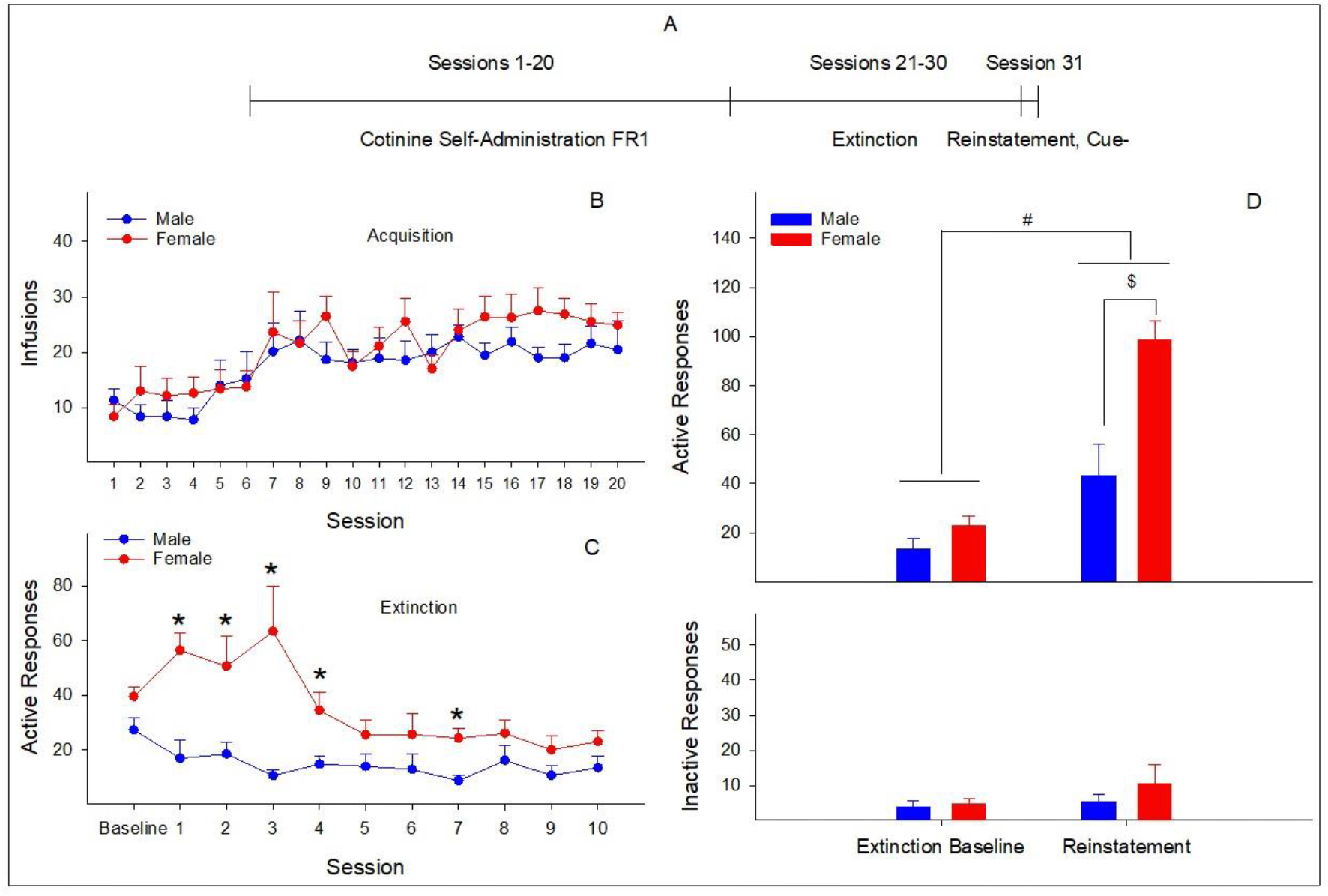
Cotinine (0.03 mg/kg/infusion) self-administration, extinction, and cue-induced reinstatement of cotinine seeking in male and female rats. (A) The timeline of self-administration, extinction, and reinstatement phases. (B) Number of cotinine infusions across self-administration sessions. (C) Active lever responses during extinction sessions. Baseline: average active responses during the last three self-administration sessions. *: *p* < 0.05, significantly greater than responses in male rats. (D) Lever responses during cue-induced reinstatement of cotinine seeking. Extinction baseline: average level responses during the last three extinction sessions. #: *p* < 0.05, significantly different between extinction baseline and reinstatement. $: *p* < 0.05, significantly different between male and female rats.

### Cotinine-induced reinstatement of drug-seeking behavior

There was significant effect of session (F_19, 380_ = 7.6, *p* = 0.000), but not sex (F_1, 20_ = 0.1, *p* = 0.70) or sex x session interaction (F_19, 380_ = 0.9, *p* = 0.53) during cotinine self-administration (Fig. 2B). Cotinine infusions increased overtime in both male and female rats. During extinction (Fig. 2C), there were significant effects of session (F_10, 200_ = 12, *p* = 0.000) and sex x session interaction (F_10, 200_ = 5.8, *p* = 0.000), but no significant effect of sex (F_1, 20_ = 0.7, *p* = 0.43). Both male and female rats reduced lever responses over time, with male rats showing greater active responses during the first two extinction sessions than female rats. During reinstatement, there were significant effects of session (F_2, 40_ = 9.9, *p* = 0.000) and sex (F_1, 20_ = 7.1, *p* = 0.014), but no significant effect of sex x session interaction (F_2, 40_ = 1.5, *p* = 0.24) on active responses (Fig. 2D, upper panel). Cotinine at both doses increased active responses regardless of sex compared to the vehicle treatment. The significant sex effect appeared to be mainly driven by greater responses in female than male rats following the 2.0 mg/kg cotinine treatment. On the other hand, cotinine did not alter inactive responses (Fig. 2D, bottom panel) with no significant effect of session (F_2, 40_ = 1.3, *p* = 0.3), sex (F_1, 20_ = 0.1, *p* = 0.31), or sex x session interaction (F_2, 40_ = 1.7, *p* = 0.20).

**Figure 2.**
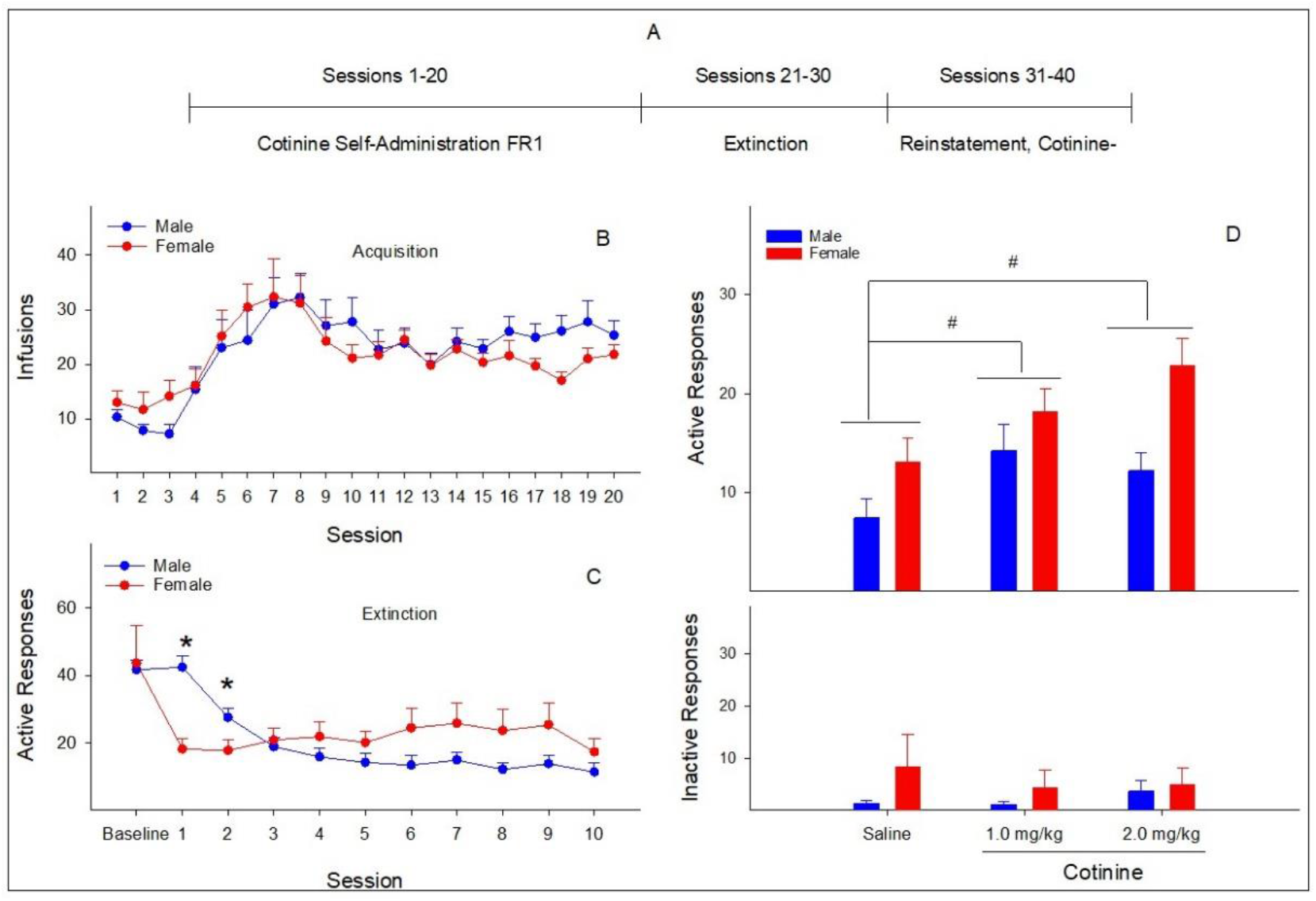
Cotinine (0.03 mg/kg/infusion) self-administration, extinction, and cotinine-induced reinstatement of cotinine seeking in male and female rats. (A) The timeline of self-administration, extinction, and reinstatement phases. (B) Number of cotinine infusions across self-administration sessions. (C) Active lever responses during extinction sessions. Baseline: average active responses during the last three self-administration sessions. *: *p* < 0.05, significantly greater responses in male than female rats. (D) Lever responses during cotinine-induced reinstatement of cotinine seeking. #: *p* < 0.05, significantly different between saline and cotinine treatments.

### Yohimbine-induced reinstatement of drug-seeking behavior

During self-administration (Fig. 3B), there was significant effects of session (F_19, 304_ = 4.5, *p* = 0.000) and sex (F_1, 16_ = 5.5, *p* = 0.032), but no significant effect of sex x session interaction (F_19, 304_ = 1.4, *p* = 0.11). Both male and female rats increased cotinine infusions overtime with female rats showing greater overall numbers of infusions than male rats. During extinction (Fig. 3C), there was significant effect of session (F_10, 160_ = 7.5, *p* = 0.000), but not sex (F_1, 16_ = 2.9, *p* = 0.11), or sex x session interaction (F_10, 160_ = 0.7, *p* = 0.69) on active lever responses. Both male and female rats reduced lever responses over time. During reinstatement, there was significant effect of session (F_2, 32_ = 25.5, *p* = 0.000), but not sex (F_1, 16_ = 1.3, *p* = 0.27) or sex x session interaction (F_2, 32_ = 1.6, *p* = 0.22) on active responses (Fig. 3D, upper panel). Yohimbine at both doses increased active responses regardless of sex compared to the vehicle treatment. For inactive lever (Fig. 3D, bottom panel), there was no significant effect of session (F_2, 32_ = 3.3, *p* = 0.05), sex (F_1, 16_ = 4.4, p = 0.051), or sex x session interaction (F_2, 32_ = 0.3, *p* = 0.75). Yohimbine did not alter inactive responses.

**Figure 3.**
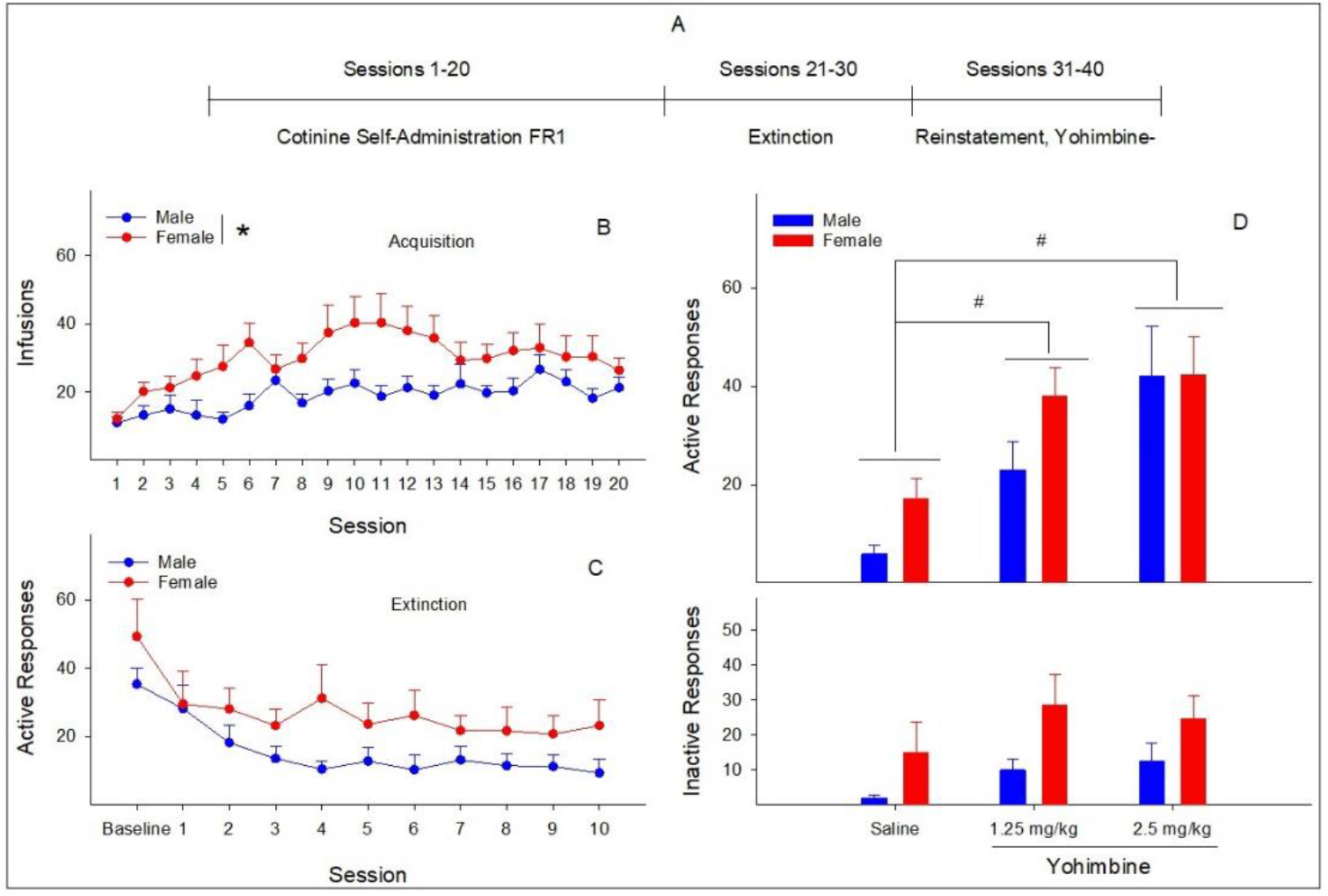
Cotinine (0.03 mg/kg/infusion) self-administration, extinction, and yohimbine-induced reinstatement of cotinine seeking in male and female rats. (A) The timeline of self-administration, extinction, and reinstatement phases. (B) Number of cotinine infusions across self-administration sessions. *: *p* < 0.05, significantly different between male and female rats. (C) Active lever responses during extinction sessions. Baseline: average active responses during the last three self-administration sessions. (D) Lever responses during yohimbine-induced reinstatement of cotinine seeking. #: *p* < 0.05, significantly different between saline and yohimbine treatments.

### Incubation of drug craving in rats

During self-administration (Fig. 4B), there was significant effects of session (F_13, 195_ = 3.5, *p* = 0.000), but not sex (F_1, 15_ = 0.3, *p* = 0.59) or sex x session interaction (F_13, 195_ = 1.0, *p* = 0.43). Both male and female rats increased number of infusions overtime. For incubation of drug craving, there was significant effect of session (F_1, 15_ = 13.3, *p* = 0.002), but no significant effect of sex (F_1, 15_ = 1.7, *p* = 0.21) or sex x session interaction (F_1, 15_ = 0.5, *p* = 0.83) on active responses (Fig. 4C, upper panel). Returning rats to the drug-associated context on withdrawal day 18 increased active responses regardless of sex compared to the responses during the withdrawal day 1. For inactive lever (Fig. 4C, bottom panel), there was no significant effect of session (F_1, 15_ = 2.7, *p* = 0.12), sex (F_1, 15_ = 0.4, *p* = 0.54), or sex x session interaction (F_1, 15_ = 0.4, *p* = 0.54). The prolonged withdrawal did not alter inactive responses.

**Figure 4.**
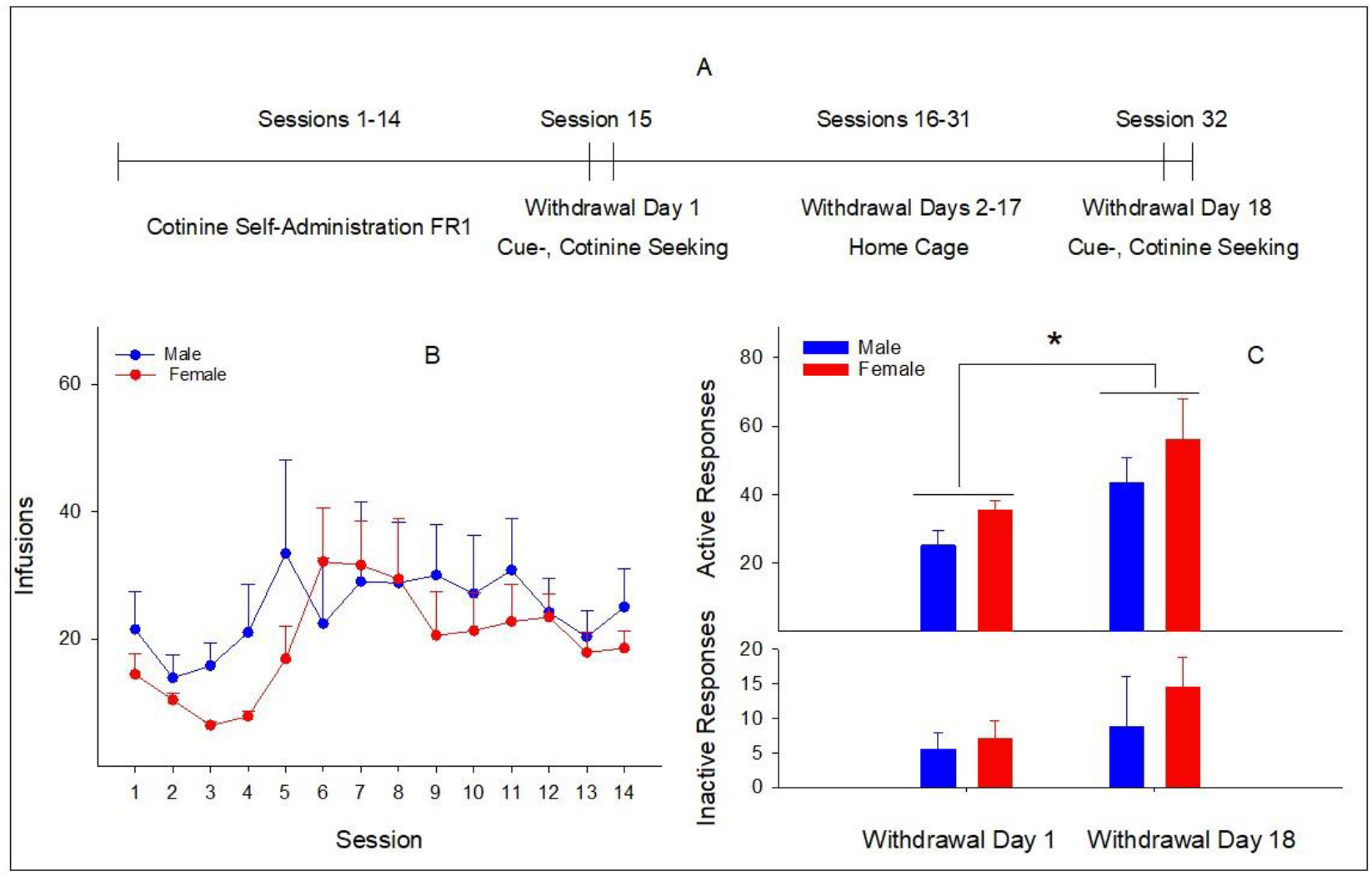
Incubation of cotinine craving in male and female rats. (A) The timeline of cotinine (0.03 mg/kg/infusion) self-administration, withdrawal, and cue-induced cotinine seeking during withdrawal. (B) Numbers of infusions during cotinine self-administration. (C) Lever responses during cue-induced cotinine seeking on withdrawal days 1 and 18. *: *p* < 0.05, significantly different between withdrawal day 18 and withdrawal day 1.

### Sex differences in cotinine self-administration and extinction

In experiments involving the reinstatement of cotinine seeking, sex differences in self-administration and/or extinction were observed in some but not other experiments. Therefore, this analysis determined potential overall sex differences in self-administration and/or extinction with data collapsed from all reinstatement experiments. Significant sex differences were found in the number of infusions during self-administration sessions (Fig. 5A, F_1, 56_ = 4.8, *p* = 0.033), and in active responses during extinction sessions (Fig. 5B, F_1, 55_ = 12.3, *p* = 0.001). Compared to male rats, female rats exhibited greater overall numbers of infusions during self-administration sessions, and showed greater resistance to extinction of active responses during extinction sessions.

**Figure 5.**
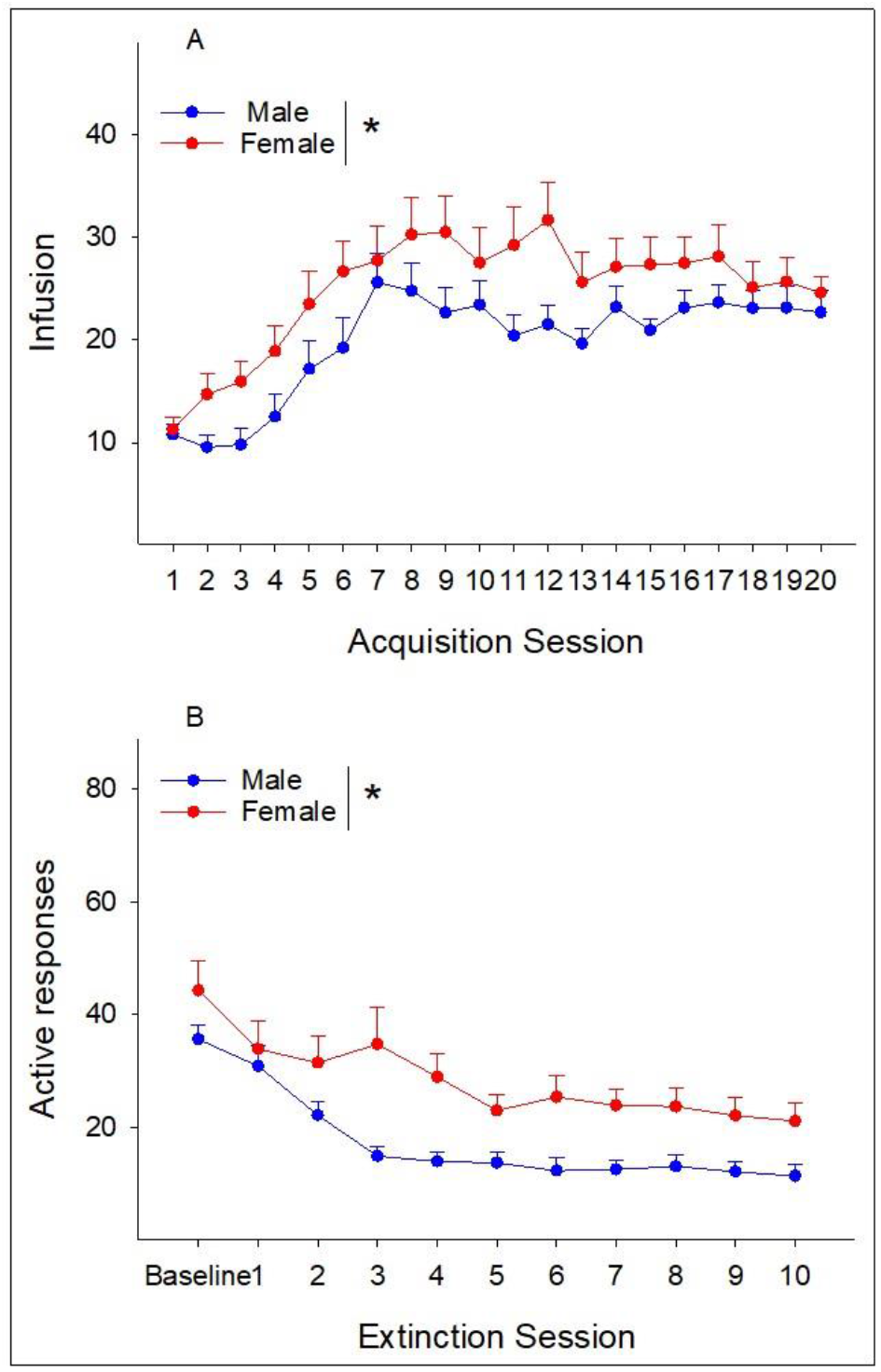
Number of cotinine infusions during self-administration (A) and active responses during extinction (B) from male and female rats included in all reinstatement experiments. *: *p* < 0.05, significantly different between male and female rats.

## Discussion

Relapse to smoking is a defining feature of human smokers and a great challenge of cessation management. A better understanding of mechanisms underlying relapse is critical to developing therapeutic strategy combating smoking. The current results support our hypothesis that cotinine, the major metabolite of nicotine, also induces relapse-like behaviors in rats as manifested by elevated cotinine-seeking behaviors in both relapse models. In the reinstatement model, contingent presentation of cues associated with cotinine self-administration, priming injection of cotinine, and the administration of the pharmacological stressor yohimbine, a α2 adrenergic receptor antagonist, augmented responses on the level previously associated with cotinine self-administration, but not responses on an inactive level. In the incubation model, the presentation of cues associated with cotinine self-administration induced greater active responses on withdrawal day 18 than on withdrawal day 1. Prior to reinstatement tests, female rats displayed greater cotinine infusions than male rats during self-administration sessions, and were more resistant to extinction of cotinine-maintained responses during extinction sessions. In addition, female rats exhibited greater cotinine-seeking behavior in cue-induced reinstatement than male rats. In contrast, there was no difference in cotinine-seeking behaviors in other relapse conditions. These findings indicate that 1) a history of cotinine self-administration can result in relapse-like cotinine-seeking behaviors, and 2) sex plays an important role in cotinine self-administration, extinction, and reinstatement. These findings suggest that cotinine may be a novel mechanism contributing to relapse to nicotine seeking.

Conditioned cues induced robust relapse to cotinine seeking in both reinstatement and the incubation of craving models (Fig. 1 & Fig. 4). These results are in agreement with previous findings indicating cue-induced nicotine seeking in the reinstatement model (Forget et al 2009, Liu et al 2008) and the incubation model (Funk et al 2016, Markou et al 2016). In the reinstatement model, female rats displayed greater cue-induced reinstatement of cotinine seeking than male rats (Fig. 1). Only a few studies examined potential sex differences in cue-induced reinstatement of nicotine seeking. One study reported greater nicotine seeking in male than female rats (Goenaga et al 2019), but another study did not find significant difference between male and female rats (Feltenstein et al 2012). These studies and our findings suggest complex sex effects on and different mechanisms involved in cue-induced reinstatement of nicotine and cotinine seeking. On the other hand, there was no significant difference in cue-induced cotinine seeking between male and female rats in the incubation model (Fig. 4). Currently, there is a lack of information on potential sex effect on cue-induced nicotine seeking in the incubation model, which will be interesting for future research.

It is interesting that the sex difference in cue-induced cotinine seeking was only observed in the reinstatement model, but not in the incubation model, which suggests a model-dependent sex effect. One major difference between these two models is the inclusion of an extinction training in the reinstatement model. Note that female rats in the reinstatement experiment also exhibited higher active responses than male rats during early extinction sessions (Fig. 1). This positive association between the greater resistance to extinction and the greater cue-induced seeking in female rats suggest that the extinction training may play an important role in manifesting the sex effect on cue-induced cotinine seeking. A previous study demonstrated that different magnitude of relapse to nicotine seeking was found between the reinstatement and incubation models (Markou et al 2016). Although no sex difference was investigated in the Markou et al study, this study and our findings suggest that the extinction training can produce various model-dependent effects on cue-induced seeking. In addition, the Markou et al study was conducted in male rats and reported less robust nicotine seeking in the reinstatement model than in the incubation model. Our study revealed no difference in male rats in cotinine seeking between the two models, but found elevated cotinine seeking in female rats in the reinstatement model compared to the incubation model (Fig. 1 & 4). Future research will be needed to address potential sex effects on cue-induced cotinine seeking in the incubation model.

The pharmacological stressor, yohimbine, reinstated cotinine seeking (Fig. 3), which is line with previous studies demonstrating yohimbine-induced reinstatement of nicotine seeking (Feltenstein et al 2012, Yuan et al 2017). In addition, there was no sex effect on yohimbine-induced cotinine seeking (Fig. 3). Similarly, a previous study revealed no significant difference between male and female rats in yohimbine-induced nicotine seeking (Feltenstein et al 2012). The qualitative similarity between nicotine and cotinine in yohimbine-reinstated seeking suggests that stress-induced reinstatement of nicotine and cotinine seeking may share some common mechanism(s). Yohimbine is a widely used pharmacological stressor in studying stress-reinstated drug seeking, and has been successfully employed to reinstate seeking for other drugs of abuse, including psychostimulants, opioids, and alcohol (Mantsch et al 2016). Although yohimbine is a reliable and robust stressor, certain weakness associated with yohimbine use has been recognized. Yohimbine was shown to enhance both inactive and active responses in some studies, which suggest a non-specific increase of general activity (Grella et al 2014). Our results indicate a strong trend for increased inactive responses following yohimbine challenge (*p* = 0.05, Fig. 3). Given this potential disadvantage, it will be of interest for future studies to test other stressors, e.g., footshock, on the reinstatement of cotinine seeking.

Acute exposure to a priming injection of cotinine also induced reinstatement of cotinine seeking (Fig. 2). Cotinine only increased active responses, but not inactive responses, suggesting that the effects of cotinine may not be due to elevation of general activity. Indeed, previous studies showed that cotinine in the range of 1-2.0 mg/kg did not alter locomotor activity in rats (Wang et al 2020b, Wiley et al 2015). Our finding is consistent with numerous studies reporting nicotine-reinstated nicotine seeking in rats (O’Connor et al 2010, Shaham et al 1997, Stoker & Markou 2015). The effects of cotinine appeared to be less robust than those of conditioned cues and yohimbine, which suggests that rats with a history of cotinine self-administration may have heightened responsiveness to cue- and stress-induced relapse than drug-induced reinstatement. These results suggest that both cotinine and nicotine can contribute to the reinstatement of drug seeking behavior, possibly via some common mechanism(s). This hypothesis can be further tested with studies examining effects of cotinine priming injections on drug seeking in subjects with a history of nicotine self-administration, and vice versa.

Significant sex differences were also observed in cotinine self-administration and extinction in several subsets of rats in the current study. For example, one subset of rats showed more cotinine infusions in female than male rats during self-administration sessions (Fig. 3). Female rats were more resistant to extinction than male rats in one subset of rats (Fig. 1), but were less resistant to extinction than male rats in another subset of rats (Fig. 2). These sporadic findings prompted an overall analysis of all rats undergoing both self-administration and extinction training; the analysis revealed that female rats showed greater self-administration and less extinction than male rats (Fig. 5). It is noted that these sex differences were only evident in some but not all individual experiments in the current study and a previous study (Ding et al 2021). It is possible that these sex differences are subtle and highly variable among individual studies; therefore, the manifestation of significant sex effects may depend on sample size and/or animals tested.

Interestingly, similar patterns of sex differences have been reported in studies of nicotine self-administration. It is generally believed that females are more vulnerable and responsive to nicotine reinforcement and use (O’Dell & Torres 2014). However, this sex difference was only observed in some studies (Donny et al 2000, Rezvani et al 2008, Sanchez et al 2014), but not others (Goenaga et al 2019, Li et al 2012, Rezvani et al 2018). Although these results are mixed, a recent meta-analysis reviewed 20 studies published prior to 2018 that included both male and female rats for nicotine self-administration, and revealed a small, but significant sex effect (effect size at 0.18) on nicotine self-administration, i.e., female rats self-administered more nicotine than male rats (Flores et al 2019). These findings and our results suggest that cotinine may contribute to sex differences in nicotine self-administration. On the other hand, potential sex differences in extinction of nicotine-reinforced operant responding have not been extensively studied. Several studies reported that male and female rats followed similar extinction curves after nicotine self-administration, suggesting no significant sex difference in extinction (Feltenstein et al 2012, Goenaga et al 2019, Swalve et al 2016). These findings suggest that extinction of nicotine- and cotinine-reinforced responding may involve different mechanisms. This also highlights the need for more research on potential sex differences in extinction of nicotine-reinforced behavior.

The sex differences in cotinine self-administration, extinction, and reinstatement are consistent with certain findings from human studies. For example, women, in general, tend to be more sensitive and susceptible to tobacco smoking, display higher rates of responding for smoking-related cues, and are less successful in quitting smoking than men (reviewed in (O’Dell & Torres 2014, Perkins 2001). Our findings suggest that sex differences in cotinine-reinforced responding may contribute to sex differences in human smokers. Therefore, elucidation of mechanisms underlying cotinine self-administration, extinction, and reinstatement in both male and female rats may provide better understanding of smoking, abstinence, and relapse in men and women smokers.

Although the current study demonstrated significant sex effects on cotinine-reinforced behaviors, it is noted that cotinine was only tested at one dose, i.e., 0.03 mg/kg/infusion. Our previous study (Ding et al 2021) indicated that cotinine was self-administered by rats in a dose-dependent manner with doses in the range of 0.0075-0.06 mg/kg/infusion. Cotinine at the dose of 0.03 mg/kg/infusion induced reliable self-administration and produced blood cotinine levels (∼ 450 ng/ml) close to those typically obtained in habitual smokers (Hukkanen et al 2005). Therefore, this optimal dose was selected for testing in this study and findings may be of good translational potentials. Another limitation is that estrous cycle was not monitored in female rats. Several studies indicated that estrous cycle did not have a major impact on nicotine self-administration or reinstatement of nicotine seeking (Donny et al 2000, Feltenstein et al 2012, Goenaga et al 2019, Rezvani et al 2008). Although these studies may suggest a general lack of effect from estrous cycle on cotinine-reinforced behaviors, this notion should be tested in the future.

In summary, the current study demonstrated robust relapse to cotinine-seeking behaviors in two well-established relapse models. Significant sex effects were also revealed that female rats exhibited greater infusions of cotinine during self-administration sessions, greater resistance to extinction of cotinine-maintained responding during extinction sessions, and greater cue-induced reinstatement of cotinine seeking. These results suggest that cotinine may play a role in relapse to nicotine seeking. In addition, these findings provide feasible models for interrogation of potential mechanisms underlying relapse to cotinine seeking. Such novel mechanisms will significantly increase our understanding of relapse to nicotine and smoking, shedding light on strategy development for aiding in smoking cessation.

## Acknowledgements

This study was supported in part by NIH grants R01DA044242 (Ding) from the US National Institute of Drug Abuse. The authors declare no conflicts of interest. The content of this manuscript is solely the responsibility of the authors and does not necessarily represent the official views of the NIH.

## Authorship Contributions

Ding participated in research design, data collection and analysis, and manuscript preparation. Tan and Neslund participated in data collection and analysis, and manuscript review.

## References

Abood LG, Grassi S, Costanza M. 1983. Binding of optically pure (--)-[3H]nicotine to rat brain membranes. FEBS Lett 157: 147–49

Anderson DJ, Arneric SP. 1994. Nicotinic recptor binding of [3H]cytisine, [3H]nicotine and [3H]methylcarbamylcholine in rat brain. Eur J Pharmacol 253: 261–67

Aubin HJ, Luquiens A, Berlin I. 2014. Pharmacotherapy for smoking cessation: pharmacological principles and clinical practice. Br J Clin Pharmacol 77: 324–36

Benowitz NL. 2010. Nicotine addiction. N Engl J Med 362: 2295–303

Benowitz NL, Jacob PI. 1994. Metabolism of nicotine to cotinine studies by a dual stable isotope method. Clin Pharmacol Ther 56: 483–93

Benowitz NL, Kuyt F, Jacob PI, Jones RT, Osman AL. 1983. Cotinine disposition and effects. Clin Pharmacol Ther 34: 604–11

Berg SA, Sentir AM, Cooley BS, Engleman EA, Chambers RA. 2014. Nicotine is more addictive, not more cognitively therapeutic in a neurodevelopment model of schizophrenia produced by neonatal ventral hippocampus lesions. Addict Biol 19: 1020–31

Boiangiu RS, Mihasan M, Gorgan DL, Stache BA, Petre BA, Hritcu L. 2020. Cotinine and 6-hydroxy-L-nicotine reverses memory deficits and reduces oxidative stress in Aβ(25-35)-induced rat model of Alzheimer’s disease. Antioxidants (Basel) 9: 768

Buczek Y, Lê AD, Wang A, Stewart J, Shaham Y. 1999. Stress reinstates nicotine seeking but not sucrose solution seeking in rats. Psychopharmacology 144: 183–88

Chiamulera C, Borgo C, Falchetto S, Valerio E, Tessari M. 1996. Nicotine reinstatement of nicotine self-administration after long-term extinction. Psychopharmacology 127: 102–07

Clemens KJ, Caille S, Stinus L, Cador M. 2009. The addition of five minor tobacco alkaloids increases nicotine-induced hyperactivity, sensitization and intravenous self-administration in rats. Int J Neuropsychopharmacol 12: 1355–66

Creamer MR, Wang TW, Babb S, Cullen KA, Day H, et al. 2019. Tobacco product use and cessation indicators among adults-United States, 2018. MMWR Morb Mortal Wkly Rep 68: 1013–19

Dar MS, Li C, Bowman ER. 1993. Central behavioral interactions between ethanol, (-)-nicotine, and (-)-cotinine in mice. Brain Res Bull 32: 23–28

Ding ZM, Gao Y, Sentir A, Tan X. 2021. Self-administration of cotinine in Wistar rats: comparisons to nicotine. J Pharmacol Exp Ther 376: 338–47

Doherty K, Kinnunen T, Militello FS, Garvey AJ. 1995. Urges to smoke during the first month of abstinence: relationship to relapse and predictors. Psychopharmacology 119: 171–8

Donny EC, Caggiula AR, Rowell PP, Gharib MA, Maldovan V, et al. 2000. Nicotine self-administration in rats: estrous cycle effects, sex differences and nicotinic receptor binding. Psychopharmacology 151: 392–405

Dwoskin LP, Teng L, Buxton ST, Crooks PA. 1999. (S)-(-)-Cotinine, the major brain metabolite of nicotine, stimulates nicotinic receptors to evoke [^3^H]dopamine release from rat striatal slices in a calcium-dependent manner. J Pharmacol Exp Ther 288: 905–11

Echeverria V, Zeitlin R, Burgess S, Patel S, Barman A, et al. 2011. Cotinine reduces amyloid-beta aggregation and improves memory in Alzheimer’s disease mice. J Alzheimers Dis 24: 817–35

Feltenstein MW, Ghee SM, See RE. 2012. Nicotine self-administration and reinstatement of nicotine-seeking in male and female rats. Drug Alcohol Depend 121: 240–6

Flores RJ, Uribe KP, Swalve N, O’Dell LE. 2019. Sex differences in nicotine intravenous self-administration: A meta-analytic review. Physiol Behav 203: 42–50

Forget B, Coen KM, Le Foll B. 2009. Inhibition of fatty acid amide hydrolase reduces reinstatement of nicotine seeking but not break point for nicotine self-administration--comparison with CB1 receptor blockade. Psychopharmacology 205: 613–24

Funk D, Coen K, Tamadon S, Hope BT, Shaham Y, Le AD. 2016. Role of Central Amygdala Neuronal Ensembles in Incubation of Nicotine Craving. J Neurosci 36: 8612–23

Goenaga J, Powell GL, Leyrer-Jackson JM, Pina J, Phan S, et al. 2019. N-acetylcysteine yields sex-specific efficacy for cue-induced reinstatement of nicotine seeking. Addict Biol

Goldberg SR, Risner ME, Stoleman IP, Reavill C, Garcha HS. 1989. Nicotine and some related compounds: effects on schedule-controlled behavior and discriminative properties in rats. Psychopharmacology 97: 295–302

Grella SL, Funk D, Coen K, Li Z, Le AD. 2014. Role of the kappa-opioid receptor system in stress-induced reinstatement of nicotine seeking in rats. Behav Brain Res 265: 188–97

Hatsukami D, Pentel PR, Jensen J, Nelson D, Allen SS, et al. 1998. Cotinine: effects with and without nicotine. Psychopharmacology 135: 141–50

Hughes JR, Keely J, Naud S. 2004. Shape of the relapse curve and long-term abstinence among untreated smokers. Addiction 99: 29–38

Hukkanen J, Jacob PI, Benowitz NL. 2005. Metabolism and disposition kinetics of nicotine. Pharmacol Rev 57: 79–115

Keenan RM, Hatsukami DK, Pentel PR, Thompson TN, Grillo M. 1994. Pharmacodynamic effects of cotinine in abstinent cigarette smokers. Clin Pharmacol Ther 55: 581–90

Li S, Zou S, Coen K, Funk D, Shram MJ, L. AD. 2012. Sex differences in yohimbine-induced increases in the reinforcing efficacy of nicotine in adolescent rats. Addict Biol 19: 156–64

Liu X, Caggiula AR, Palmatier MI, Donny EC, Sved AF. 2008. Cue-induced reinstatement of nicotine-seeking behavior in rats: effect of bupropion, persistence over repeated tests, and its dependence on training dose. Psychopharmacology 196: 365–75

Mantsch JR, Baker DA, Funk D, Le AD, Shaham Y. 2016. Stress-Induced Reinstatement of Drug Seeking: 20 Years of Progress. Neuropsychopharmacology 41: 335–56

Markou A, Li J, Tse K, Li X. 2016. Cue-induced nicotine-seeking behavior after withdrawal with or without extinction in rats. Addict Biol 23: 111–9

National Research Council. 2011. Guide for the Care and Use of Laboratory Animals. Washington, D.C.: National Academies Press.

O’Connor EC, Parker D, Rollema H, Mead AN. 2010. The α4β2 nicotinic acetylcholine-receptor partial agonist varenicline inhibits both nicotine self-administration following repeated dosing and reinstatement of nicotine seeking in rats. Psychopharmacology 208: 365–76

O’Dell LE, Torres OV. 2014. A mechanistic hypothesis of the factors that enhance vulnerability to nicotine use in females. Neuropharmacology 76: 566–80

Perkins KA. 2001. Smoking Cessation in Women: Specifial considerations. CNS Drugs 15: 391–411

Pickens CL, Airavaara M, Theberge F, Fanous S, Hope BT, Shaham Y. 2011. Neurobiology of the incubation of drug craving. Trends Neurosci 34: 411–20

Plaza-Zabala A, Martin-Garcia E, de Lecea L, Maldonado R, Berrendero F. 2010. Hypocretins regulate the anxiogenic-like effects of nicotine and induce reinstatement of nicotine-seeking behavior. J Neurosci 30: 2300–10

Prochaska JJ, Benowitz NL. 2016. The Past, Present, and Future of Nicotine Addiction Therapy. Annu Rev Med 67: 467–86

Rezvani AH, Eddins D, Slade S, Hampton DS, Christopher NC, et al. 2008. Neonatal 6-hydroxydopamine lesions of the frontal cortex in rats: persisting effects on locomotor activity, learning and nicotine self-administration. Neuroscience 154: 885–97

Rezvani AH, Tizabi Y, Slade S, Getachew B, Levin ED. 2018. Sub-anesthetic doses of ketamine attenuate nicotine self-administration in rats. Neurosci Lett 668: 98–102

Risner ME, Goldberg SR, Prada JA, Cone EJ. 1985. Effects of nicotine, cocaine and some of their metabolites on schedule-controlled responding by beagle dogs and squirrel monkeys. J Pharmacol Exp Ther 234: 113–19

Rosecrans JA, Chance WT. 1977. Cholinergic and non-cholinergic aspects of the discriminative stimulus properties of nicotine In Discriminative stimulus properties of drugs, ed. H Lal, pp. 155–86. New York: Plenum Press

Rosen LJ, Galili T, Kott J, Goodman M, Freedman LS. 2018. Diminishing benefit of smoking cessation medications during the first year: a meta-analysis of randomized controlled trials. Addiction 113: 805–16

Sanchez V, Moore CF, Brunzell DH, Lynch WJ. 2014. Sex differences in the effect of wheel running on subsequent nicotine-seeking in a rat adolescent-onset self-administration model. Psychopharmacology 231: 1753–62

Schroff KC, Lovich P, Schmitz O, Aschhoff S, Richter E, Remien J. 2000. Effects of cotinine at cholinergic nicotinic receptors of the sympathetic superior cervical ganglion of the mouse. Toxicology 144: 99–105

Shaham Y, Adamson LK, Grocki S, Corrigall WA. 1997. Reinstatement and spontaneous recovery of nicotine seeking in rats. Psychopharmacology 130: 396–403

Shaham Y, Shalev U, Lu L, De Wit H, Stewart J. 2003. The reinstatement model of drug relapse: history, methodology and major findings. Psychopharmacology 168: 3–20

Stoker AK, Markou A. 2015. Neurobiological Bases of Cue-and Nicotine-induced Reinstatement of Nicotine Seeking: Implications for the Development of Smoking Cessation Medications. Curr Top Behav Neurosci 24: 125–54

Substance Abuse and Mental Health Services Administration (SAMHSA). 2019. Key Substance Use and Mental Health Indicators in the United States: Results from the 2018 National Survey on Drug Use and Health. HHS Publication No. PEP19-5068, NSDUH Series H-54 Rockville MD: Center for Behavioral Health Statistics and Quality, Substance Abuse and Mental Health Services Administration

Swalve N, Smethells JR, Carroll ME. 2016. Sex differences in attenuation of nicotine reinstatement after individual and combined treatments of progesterone and varenicline. Behav Brain Res 308: 46–52

Takada K, Swedberg MD, Goldberg SR, Katz JL. 1989. Discriminative stimulus effects of intravenous l-nicotine and nicotine analogs or metabolites in squirrel monkeys. Psychopharmacology 99: 208–12

Tan X, Vrana K, Ding Z. 2021. Cotinine: Pharmacologically active metabolite of nicotine and neural mechanisms for its actions. Front Behav Neurosci 15: 758252

Terry Jr AV, Hernandez CM, Hohnadel EJ, Bouchard KP, Buccafusco JJ. 2005. Cotinine, a neuroactive metabolite of nicotine: poetential for treating disorders of impaired cognition. CNS Drug Rev 11: 229–52

U.S. Department of Health and Human Services (HHS). 2014. The health consequences of smoking--50 years of progress. Atlanta, GA: U.S. Department of Health and Human Services, CDC.

Venniro M, Caprioli D, Shaham Y. 2016. Animal models of drug relapse and craving: From drug priming-induced reinstatement to incubation of craving after voluntary abstinence. Prog Brain Res 224: 25–52

Wang TW, Neff LJ, Park-Lee E, Ren C, Cullen KA, King BA. 2020a. E-cigarette use among middle and high school students-United States, 2020. MMWR Morb Mortal Wkly Rep 69: 1310–12

Wang Y, Wan B, Huang J, Clarke PBS. 2020b. Effects of nicotine, nornicotine and cotinine, alone or in combination, on locomotor activity and ultrasonic vocalization emission in adult rats. Psychopharmacology 237: 2809–22

Ward SJ, Rosenberg M, Dykstra LA, Walker EA. 2009. The CB1 antagonist rimonabant (SR141716) blocks cue-induced reinstatement of cocaine seeking and other context and extinction phenomena predictive of relapse. Drug Alcohol Depend 105: 248–55

Wiley JL, Marusich JA, Thomas BF, Jackson KJ. 2015. Determination of behaviorally effective tobacco constituent doses in rats. Nicotine Tob Res 17: 368–71

Yuan M, Malagon AM, Yasuda D, Belluzzi JD, Leslie FM, Zaveri NT. 2017. The α3β4 nAChR partial agonist AT-1001 attenuates stress-induced reinstatement of nicotine seeking in a rat model of relapse and induces minimal withdrawal in dependent rats. Behav Brain Res 333: 251–57

